# The causal arrows ̶ from genotype, environment and management to plant phenotype ̶ are double headed

**DOI:** 10.1101/2024.04.08.588646

**Authors:** Victor O Sadras, Peter T Hayman

**Affiliations:** South Australian Research and Development Institute; School of Agriculture, Food and Wine, The University of Adelaide; College of Science and Engineering, Flinders University, Australia

**Keywords:** behaviour, context, development, downward causation, farmer, mutation, niche construction, risk, rotation, teleonomy, water

## Abstract

Unidirectional, cause-and-effect arrows are drawn from genotype (G), environment (E), and agronomic management (M) to the plant phenotype in crop stands. Here we focus on the overlooked bidirectionality of these arrows. The phenotype-to-genotype arrow includes increased mutation rates in stressed phenotypes, relative to basal rates. From a developmental viewpoint, the phenotype modulates gene expression returning multiple cellular phenotypes with a common genome. From a computational viewpoint, the phenotype influences gene expression in a process of downward causation. The phenotype-to-environment arrow is captured in the process of niche construction, which spans from persistent and global (*e.g*., photosynthetic archaea and cyanobacteria that emerged ∼3.4 billion years ago *created* the oxygen-rich atmosphere that *enabled* the evolution of aerobic organisms and eukaryotes) to transient and local (*e.g.*, lucerne tap root constructs soil biopores that influence the root phenotype of the following wheat crop). Research on crop rotations illustrates but is divorced of niche construction theory. The phenotype-to-management arrow involves, for example, a diseased crop that triggers fungicide treatments. Making explicit the bidirectionality of the arrows in the G **×** E **×** M model allows to connect crop improvement and agronomy with other, theoretically rich scientific fields.

**Highlight:** In the G **×** E **×** M model, the plant phenotype is not only influenced by but also influences G, E and M.

## Introduction

Organisms have two parts: the genome and the rest; the rest is the phenotype (West-Eberhard, 2003). mRNA, DNA repair enzymes, concentration of abscisic acid in maize xylem, soybean root depth, wheat yield, and the amount of anthocyanins in grapevine berries are all aspects of the phenotype. Farmers use two technologies to manipulate the phenotype of both plants in crop stands and other agronomically relevant organisms (*e.g.,* weeds, herbivores, pathogens): varieties or hybrids and agronomic practices, with a frequent synergy between improved plants and agronomy (Fischer, 2009). The phenotypic variance of a trait is usually partitioned into genetic (G) and environmental components (E) with a trait-dependent G x E interaction and residuals (Diouf *et al*., 2020; Fisher, 1919; Wright, 1920). The interaction is, for example, lower for seed weight than for seed number (Sadras, 2021), traits that are related in a hierarchy of plasticities (sensu Bradshaw, 1965). Depending on the focus of research, management (M) can be made explicit in a G **×** E **×** M model (Cooper *et al*., 2020; Hajjarpoor *et al*., 2022; Stöckle and Kemanian, 2020), or can be embedded in the environment as most practices, such as irrigation and fertilisation, seek to modify the environment. An extended G **×** E **×** M **×** S framework incorporates social factors S (Gerullis *et al*., 2023; Kholová *et al*., 2021). Unidirectional, convergent cause-and-effect arrows are drawn from genotype, environment, and management to phenotype; the bidirectionality of these arrows, the focus of this paper, is often overlooked.

In a context of system and complexity thinking in agriculture, cognitive maps have been advanced that include six motifs (Fig. 1). Of these motifs, convergent arrows representing multiple factors driving an outcome were very common (motif 2 in Fig. 1); for example, daylength and temperature, and photoperiod (*Ppd*) and vernalisation (*Vrn*) alleles converge to modulate wheat flowering time (Bloomfield *et al*., 2018). Bidirectional arrows representing mutual influences were cognitively rarer (motif 1 in Fig. 1). Here we look at the G **×** E **×** M framework with a focus on the causal relations from phenotype to genotype, *e.g.,* increased mutation rates in stressed phenotypes (arrow 1, Fig. 2); from phenotype to environment in the process of niche construction (arrow 2, Fig. 2); and from phenotype to management, *e.g.*, a diseased crop that triggers fungicide treatments (arrow 3, Fig. 2). We emphasise the living components of the plant’s environment including the crop plant influencing its neighbouring plants in the stand, and the phenotypes of farmers (Box 1) with their own sources of variation including their biophysical, economic and legal environments, all of which are unprestatable (Kauffman, 2008, 2016).

**Figure 1.**
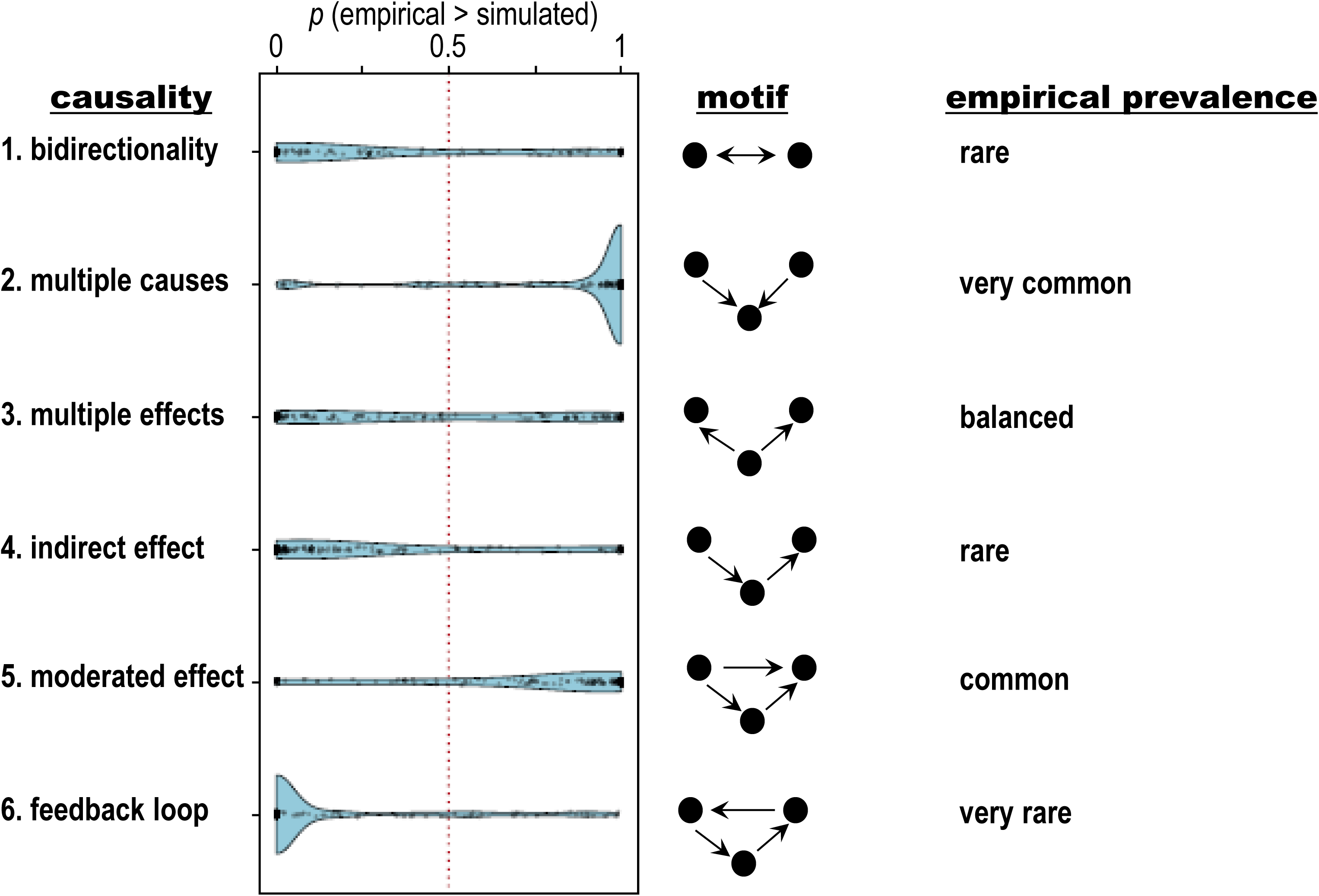
Cognitive maps of thought leaders in sustainable agriculture. The maps include six fundamental causal relationships; from top to bottom: bidirectionality, the focus of this paper; multiple causes, the default in the G x E x M model of the phenotype; multiple effects; indirect effects; moderated effect; and feedback loop. The blue graphs are the distributions of the prevalence of causal motifs in cognitive maps relative to uniform random graphs. Within each structure, each point represents one individual’s mental model; probabilities near one indicate an abundance of the structure relative to chance and the red dotted line at 0.50 indicates the expectation for each structure in a uniform random graph of the same size and density as the cognitive map. Data are from a sample of 148 experts in California, with a median experience in agriculture of 20 years; extrapolations are thus not warranted. Redrawn from Table 1 and Fig. 1 in Levy et al. (2018).

**Figure 2.**
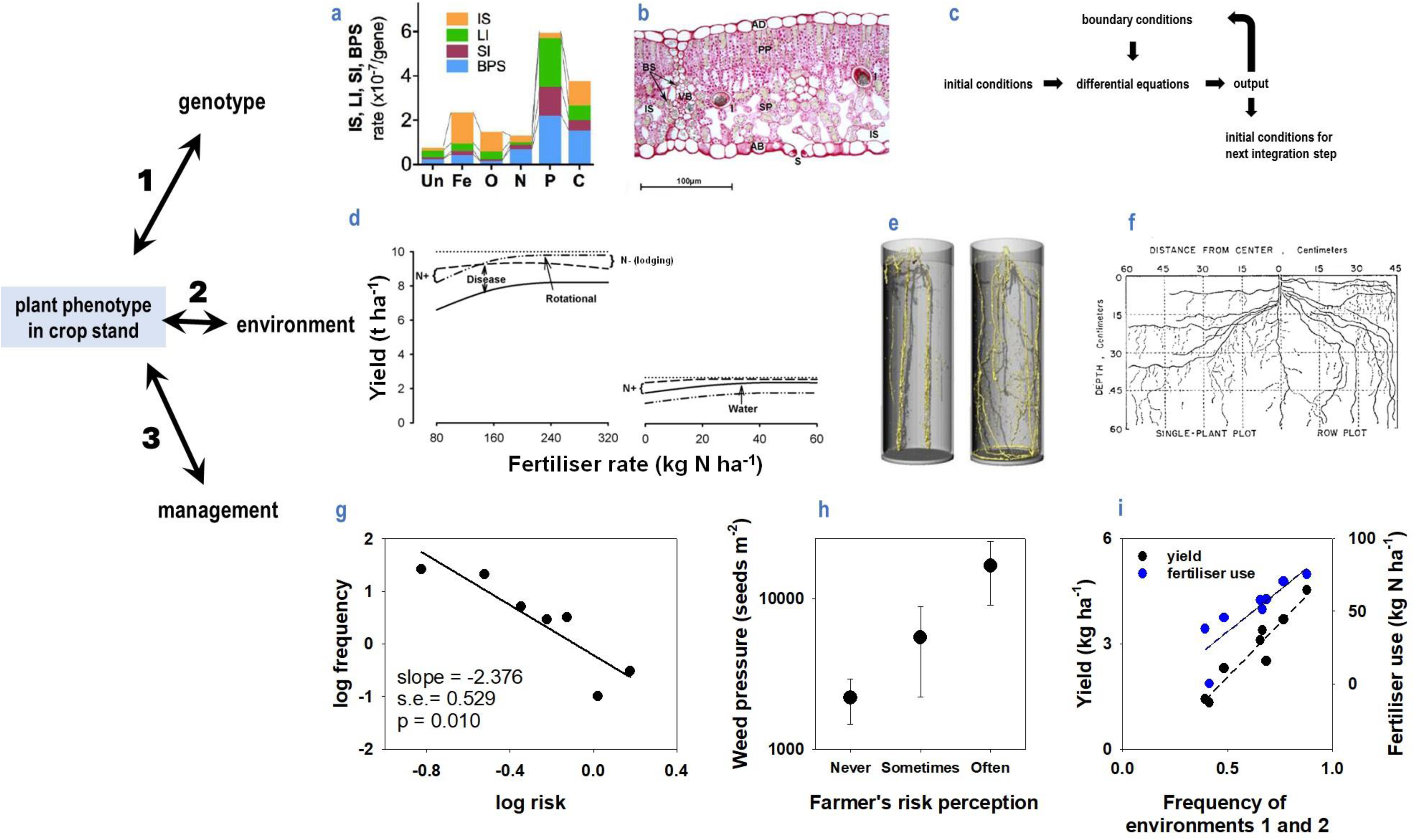
The causal relations from phenotype to genotype (arrow 1), environment (arrow 2), and management (arrow 3). **(a)** The rate of mutation increases in stressed phenotypes. Mutations include base-pair substitutions (BPS), single-base pair indels (SI), deletion and insertion indels > 1bp (LI), and insertion sequence transpositions (IS) for *Escherichia coli* in nutrient-unlimited culture (Un) and in cultures with iron (Fe), oxygen (O), nitrogen (N), phosphorus (P) and carbon limitation (C). **(b)** A single genotype returns diverse cellular phenotypes as illustrated in a leaf of Syrah featuring AB, abaxial epidermis; AD, adaxial epidermis; BS, bundle sheath; I, idioblast; PP, palisade parenchyma; S, stomata; SP, spongy parenchyma; VB, vascular bundle. **(c)** From a computational viewpoint, the arrow from genome to phenotype could be represented with differential equations that are necessary but not sufficient: the phenotype defines the initial and boundary conditions required for integration. **(d)** Yield of wheat as a function of nitrogen fertiliser rate in response to previous crop under two scenarios: left, high water availability, high agronomic input, and severe disease incidence; right, low water availability, low agronomic input, and low disease incidence. The curves show yield potential (doted), and yield of wheat after wheat (solid), after legume (dashed) and after oilseed crop (doted-dashed). **(e)** Maize roots in soil with bulk density (left) 1.8 g cm^-3^ and (right) 1.4 g cm^-3^. **(f)** Close-to-horizontal root branching in isolated soybean plants in contrast to the abrupt verticalization of roots in the presence of neighbours. **(g)** Frequency distribution of risk attitude in a sample of 313 apple growers in northern Italy, where hail and frost are major sources of risk. Risk attitude of each farmer was elicited using a lottery-choice in which subjects were confronted with a set of 50-50 gambles, including a sure outcome and several risky outcomes with linearly increasing expected payoffs and risk, measured as the standard deviation of expected payoffs. **(h)** Weed pressure in Dutch organic fields increased with farmers’ risk perception of soil structural damage associated with mechanic weed control; “never”, “sometimes” and “often” are answers to the question How often is the risk of soil structural damage a reason not to control weeds? **(i)** The doble effect of water availability on crop yield includes a direct biological component, and a component mediated by farmer’s risk attitude and input use. Seasonal water supply:demand in eight wheat-growing regions of Argentina clustered in four environment types, with environments 1 and 2 representing lack of or mild drought. Yield in commercial fields declines with lower frequency of less risky environments 1 and 2, and part of this decline associates with lower use of fertiliser. Sources: (a) Maharjan and Ferenci (2017), (b) Gago et al. (2019), (c) Noble (2012), (d) Kirkegaard et al. (2008), (e) Wendel et al. (2022), (f) Raper and Barber (1970), (g) Menapace et al. (2013), (h) Riemens et al. (2010), (i) Pellegrini et al. (2022).

## The causal arrow from phenotype to genotype: mutation rates under stress and downward causation

The causal arrow from phenotype to genotype includes two aspects. First, to strictly qualify as a causal relation, we consider changes in the phenotype that drive genotypic change with ecological and evolutionary consequences. The perspective of evolution has shifted from a process primarily associated with random mutations and natural selection to the contemporary view whereby organisms are active agents of their own genomic, phenotypic, and adaptive changes (Corning *et al*., 2023; Shapiro, 2022). The informatic metaphor has shifted from a genome as a Read-Only Memory (ROM) to a read-write (RW) data storage system subject to cellular modifications and inscriptions at three scales: cell reproduction, multicellular development, and evolutionary, and from point mutations to large-scale genome rearrangements (Shapiro, 2022). Mutation rates have traditionally been considered low, constant, and independent of the phenotype and the environment partially because most sporadic mutations are neutral or deleterious, hence the assumed adaptive value of low rates, limited only by the cost of avoidance and correction of errors (Ram and Hadany, 2019; Taddei *et al*., 1997). In this context, at least two observations justify the strict arrow from phenotype to genotype. The innate rate of error in DNA replication is typically 1 in 10,000 and is lowered to 1 in 10 billion in a vigilance process that involves a suit of unique repair enzymes (Noble and Noble, 2023); these enzymes are phenotype by definition (West-Eberhard, 2003). This provides a way for the cell to alter the DNA in a targeted process captured in the metaphor “the genes dance to the tune of the cell” (Noble and Noble, 2023). The arrow is also justified because mutation rates are higher in stressed phenotypes (Fig. 2a) as found across taxa including mammals, plants, bacteria and yeast (DeFranco, 2016; Gullickson *et al*., 2022; Maharjan and Ferenci, 2017; McClintock, 1984; Shapiro, 2022; Shewaramani *et al*., 2017). Whereas the literature on stress-induced mutagenesis usually emphasises the stress factors such as radiation, pathogens, or anaerobiosis, what matters functionally is the stressed phenotype; McClintock (1984) underscored, for example, that infection of maize plants with barley stripe mosaic virus “may traumatise cells to respond by activating potentially transposable elements”. Our current understanding of the immune system is possibly the most compelling evidence for the causal arrow from phenotype to genotype (DeFranco, 2016; Gullickson *et al*., 2022; Noble and Noble, 2023; Shapiro, 2022). Diversity of antibodies (immunoglobulins) that neutralise pathogens and their gene products is crucial for a functional immune system. This diversity stems from three processes: V(D)J recombination (<u>V</u>ariable, <u>D</u>iversity and <u>J</u>oining gene segments), class switch recombination (CSR), and somatic hypermutation (SHM), which are in turn promoted by environmental factors, chiefly the presence of antigens (Gullickson *et al*., 2022). For example, naïve B cells produce only membrane-bound antibodies Igm and Igd but naïve B cells are activated and undergo CSR that ‘fine tunes’ B cell receptors in the presence of antigens. The mutation frequency of SHM is 10^6^ higher than the basal mutation rate and conforms to the concept of intentional DNA modification that leads to high-affinity antibodies.

Mutators ̶ individuals in a population with an above-average mutation rate ̶ often arise spontaneously during evolution (Lobinska *et al*., 2023; Sane *et al*., 2023; Taddei *et al*., 1997; Tanaka *et al*., 2003). Models accounting for modifiers of the mutation rate in clonal populations showed that stable environments would select for a minimal mutation rate, but in more realistic, variable environments, populations at equilibrium could have mutation rates well above the minimum (Taddei *et al*., 1997; Tanaka *et al*., 2003). Furthermore, the adaptive superiority of mutators can also relate to an intriguingly lower frequency of deleterious mutations than in their wildtype counterparts. In an experimental comparison, a mutator strain of *Escherichia coli* created by deletion of a DNA repair gene returned a deleterious : neutral : beneficial ratio of mutations of 24 : 40 : 36 in comparison to 39 : 33 : 28 for the wild type across several environments (Sane *et al*., 2023). The mutator state could be not only genetically inherited from loss-of-function mutations in DNA repair genes (Sane *et al*., 2023), but can also arise through transgenerational epigenetic inheritance (Lobinska *et al*., 2023). Under the modelling assumptions of Lobinska et al. (2023), switching mutation rates associated with non-genetic inheritance were adaptively superior to switching rates associated with genetic inheritance. Experiments and modelling with *Saccharomyces cerevisiae* highlighted the role of both population size and migration in selecting for mutators (Raynes *et al*., 2019).

The second aspect of the arrow from phenotype to genotype is the phenotypic modulation of gene expression revealed from developmental (Levin, 2023; West-Eberhard, 2003), and physiological and computational perspectives (Noble, 2011; Noble, 2012). Wright (1920) partitioned the phenotypic variation of the piebald pattern of guinea pigs in hereditary and environmental factors, and ascribed the residual variation to an “irregularity in development”; development is not hardwired but context-sensitive and plastic (Amzallag, 2000; Levin, 2023; Schlichting, 2004). A transversal cut of a grapevine leaf highlights morphologically and functional distinct cellular phenotypes including abaxial and adaxial epidermis, bundle sheath, idioblasts, palisade parenchyma, stomata, and spongy parenchyma (Fig 2b). The same genome returns more than 30 cellular phenotypes in plants and more than 140 in vertebrates (West-Eberhard, 2003). The regulation of gene expression by abscisic acid illustrates this aspect of the phenotype-to-gene arrow (Chandler and Robertson, 1994); this is not strictly a change in genotype but is functionally relevant as the phenotype modulates itself via shifts in gene expression. From a computational viewpoint, the arrow from genome to phenotype could explain, for example, the activity of ion channels and action potentials of cell membranes (Huang *et al*., 2021; Noble, 2012), which in turn could be represented by differential equations describing the speed and the direction of the gating process on each protein (Noble, 2012). The differential equations captured in genotype-to-phenotype arrows are *necessary* but not *sufficient*; membrane and cellular traits that set the initial and boundary conditions are required to solve the biologically relevant phenotype by integration (Fig. 2c). This computational perspective converges with both the self-organising nature of specific transition phases in plant development (Amzallag, 2000) and a teleonomic (purpose oriented) model of development that proposes that to understand morphospace – the space of possible anatomical configurations that any group of cells can achieve – we need to understand not only the molecular mechanisms that are *necessary* for morphogenesis but also the information-processing dynamics that are *sufficient* for cell groups to create, repair, and reconstruct large-scale anatomical features (Levin, 2023). This is how higher scales of organisation influence lower scales in a process of downward causation, which is not mere feedback but a true cause-and-effect relation (Flack, 2017; Green, 2018; Noble, 2012).

## The causal arrow from phenotype to environment: niche construction

Deffner (2023) updated the concept of niche construction with a focus on evolution from a teleonomic perspective. During the 20^th^ century, theoretical arguments were developed for a reciprocal relationship between organism and environment, and both theory and empirical evidence have grown to return a compelling case for a causal phenotype-to-environment relation that is rarely explicit in agronomic G **×** E **×** M models (Fig. 2, arrow 2).

Niche construction is the process whereby organisms actively modify their own niche, others’ niche, or both (Odling-Smee *et al*., 2013); it spans a wide range of temporal and spatial scales. Photosynthetic archaea and cyanobacteria that emerged ∼3.4 billion years ago *created* the oxygen-rich atmosphere that *enabled* the evolution of aerobic organisms and eukaryotes 2.0-1.5 billion years ago (Baluška *et al*., 2023); this is evolutionary relevant niche construction on a geological scale.

Innovations that *enable* new innovations are at the core of the biosphere’s evolution, and this partially explains why the trajectory of the biosphere is unprestatable (Kauffman, 2008, 2016). Several species of tetranychid mites (Acari: Tetranychidae), including the two-spotted mite *Tetranychus urticae,* construct complicated three-dimensional webs on plant leaves that modify the micro-environment with consequences for the host plant, for the mites themselves, and for organisms at higher trophic levels, *e.g.,* mite predators (Oku *et al*., 2009; Roda *et al*., 2001); this is ecologically and agronomically important, transient, spatially confined niche construction.

Crop rotations are the quintessential case of niche construction in agriculture. Levantine farmers were aware of the rotational benefits of cereals and pulses in the Neolithic (Abbo and Gopher, 2022). Pliny described crop rotations in ancient Greece and Rome that are comparable to those currently used in the wheat growing regions of Australia (Sadras *et al*., 2004). A crop in the current season leaves a biological, chemical, and physical soil legacy that influences the plant phenotype of subsequent crops and other relevant phenotypes including those of weeds, pathogens, and herbivores. This soil legacy and its agronomic consequences are demonstrated in robust studies of crop rotations (Angus *et al*., 1994; Angus *et al*., 2015; Kirkegaard *et al*., 2008; Sadras *et al*., 2004). But this research is largely divorced of niche construction theory. A rare example of the interpretation of agricultural rotations in the light of niche construction theory is the study of a half-farming and half-fishing system practiced by the costal Gungokri people in southwestern Korea for five centuries since 150 BC (Lee *et al*., 2023). Rotation of crops in both wetlands and uplands sought to prevent the loss of soil nutrition and erosion from seawater; in the dry uplands, farmers mixed wheat and barley with short-lived crops such as millets, which require less nutrition, and legumes (soybean and azuki) that prevent soil erosion and add nutrition; some varieties of salinity-tolerant foxtail millet where part of the rotations (Lee *et al*., 2023).

Standard crop simulation models capture the carry-over of soil water and nitrogen but not the biological components of crop sequences; this is an important limitation (Sadras *et al*., 2004). The roots of *Brassica* spp. produce isothiocyanates that arrest the growth of *Gaeumannomyces graminis*, the fungal pathogen that causes take-all of wheat (Angus *et al*., 1994) partially contributing to the improved yield and water use efficiency of wheat after canola (*B. napus*) or mustard (*B. juncea*) compared to wheat after wheat (Angus and van Herwaarden, 2001). The total rotation effect for wheat, calculated as the change in yield of wheat after a broad-leaf break crop relative to wheat after wheat, averaged 14% in cropping environments of North America, 33% in Australia and 24% in Europe, albeit the ranges were wide including cases of negative effects (Kirkegaard *et al*., 2008). Fig. 2d illustrates the effects of rotation on wheat yield for a gradient of nitrogen fertilisation under two scenarios. In a scenario of high availability of water, high agronomic input and severe disease incidence, wheat after legume or oilseed crops typically yielded 20-30% more than wheat after wheat, and the rotation effect cannot be substituted with higher inputs (Fig. 2d, left). In a scenario of low availability of water, low agronomic input, and low disease incidence, wheat yield is largely responsive to other inputs as it primarily depends on the amount of water stored in the soil at sowing, which is generally higher following legumes than oilseeds (Fig. 2d, right). A modelling study that compared current, soybean-based cropping with alternative crop sequences including wheat and maize in the Pampas, concluded that functional crop types were more important than cropping diversity and perenniality for the profit and risk of the sequences (Videla-Mensegue *et al*., 2022). Consistently with this finding, the functional equivalence of niche constructors is more important than their identity (Deffner, 2023).

Soil biopores created by decaying roots or earthworms are another example of niche construction relevant to crops where the identity of the constructor is less important than its functionality. In soil compacted to 1.8 g cm^-3^ bulk density, maize roots only grew in pores, whereas roots grew in the matrix soil at 1.4 g cm^-3^ bulk density (Fig. 2e). Biopore construction involves a sequence of processes (Wendel *et al*., 2022). First, when available, roots and earthworms preferentially use low-penetration resistance, fine soil cracks, with roots establishing a rhizosphere and worms a drilosphere. In these spaces, nutrient cycling and microbial abundance and activity are increased compared to the bulk soil. When the root dies or the earthworm leaves the pore or dies, nutrients remain accumulated along the biopore lining and sheath. Other plant roots and earthworks can reuse the biopore reinforcing the nutrient-rich hotspot in a feedback loop (motif 6 in Fig. 1).

Plant community diversity and the phenotype of individual plants can influence the composition of their associated microbial communities, with ecological and agronomic implications. The influence of plants on their soil environment and associated microbial communities is mediated by processes such as (i) release of compounds with low molecular mass (sugars, amino acids and organic acids), polymerized sugar (*i.e.,* mucilage), root border cells and dead root cap cells; rhizodeposits account approx. for approx. 25% of the carbon allocated to the roots in cereals and grasses; (ii) release of secondary metabolites, such as antimicrobial compounds, nematicides and flavonoids, which are involved in establishing symbiosis or in warding off pathogens and pests; (iii) release and uptake of ions by roots, which can cause up to 2 units variation in soil pH; (iv) uptake of water and root respiration affecting soil moisture and oxygen pressure (Philippot *et al*., 2013). “Home-field advantage” and the co-variation between plant control of nitrification and plant preference for ammonium or nitrate are two examples of the relevance of the phenotype-to-environment causal relation in this context. It has been hypothesised that some plant species could promote the decomposition of their own litter rather than that of other plant species or genotypes returning a ‘home-field advantage’; the empirical evidence for this phenomenon is partial (Ayres *et al*., 2009; Schmitt and Perfecto, 2021). The co-variation between plant control of nitrification and plant preference for ammonium or nitrate was modelled against the hypotheses that plants with an ammonium preference would grow more biomass when inhibiting nitrification, and conversely that plants preferring nitrate would achieve higher biomass by stimulating nitrification (Ardichvili *et al*., 2024).

The model with parameters from a savanna in Ivory Coast partially supported the first hypothesis, and modelling with parameters for an intensively cultivated, short-grass prairie in the US led to the counter-intuitive combination of nitrate preference and nitrification inhibition returning higher biomass. Factors that could override the expected associations between nitrogen preference and mineralisation include quantity of nitrogen deposition in the ecosystem, leaching rates, and baseline nitrification rate (Ardichvili *et al*., 2024). Microbiology-centred research concerning agricultural impact has led to the conclusion that manipulating soil microbes could improve sustainability of cropping systems but lack of agronomic context undermines this proposition (Ryan and Graham, 2018; Ryan *et al*., 2019).

## The living components of the environment: when the environment has genes

Except where the focus is crop protection, research in plant sciences emphasises the abiotic component of the environment, chiefly resources including water and nutrients and non-resource factors such as temperature (Dalal *et al*., 2017). In a sample of 34,757 scientific papers focusing on plant stress, the abiotic : biotic ratio was 5:1 across disciplines, and it was 20 times greater in the field of ecology and 60 times greater in forestry (Dalal *et al*., 2017). The living component of the environment, generally overlooked in plant sciences using simplified experimental settings (Sadras, 2019), is important in both nature and agriculture. Darwin (1859) insisted that the relation of organism to organism is the most important of all relations, particularly against the over-rated role of adaptation to climate. The idea that “the environment of an organism mostly consists of other organisms” persists in the contemporary framework that extends the neo-Darwinian theory of evolution to account for self-organisation, symbiogenesis, teleonomy, niche construction, and genetic covariance in both heterospecific and conspecific relations (Heylighen, 2023; Wolf *et al*., 2004).

In heterospecific settings, herbivores are part of the plant’s environment and the plant is part of the herbivore’s environment; likewise, there is a reciprocal phenotype-environment relationship between rhizobia and legume plants, and between crop plants and weeds linked in co-evolutionary processes (Coba de la Peña *et al*., 2018; Guglielmini *et al*., 2007; Wolf *et al*., 2004). The two-spotted spider mite is a common secondary pest of horticultural and broadacre crops such as cotton. Owing to their size, with adults ≈ 0.5 mm and their eggs ≈ 0.1 mm, the key environment for mites is that of the boundary layer of air trapped close to the leaf surface (Wilson and Sadras, 2001). Eggs are particularly susceptible to dehydration hence the importance of the humidity of the boundary layer that varies with plant traits including transpiration rate and leaf morphological features that create regions of reduced turbulence such as high hair density, leaf folds, prominent leaf veins, and lobbed leaves (Reddall *et al*., 2011; Wilson and Sadras, 2001).

In conspecific settings such as crop stands, plant-plant interactions are primary drivers of the individual’s phenotype, and the contemporary measure of agronomic yield in annual seed crops ̶ mass of product per unit land area ̶ has favoured a communal phenotype with diminished competitive ability (Biernaskie, 2022; Cossani and Sadras, 2021; Denison, 2012; Donald, 1981; López Pereira *et al*., 2017). The zenith angle (i.e., the angle with respect to the vertical) of *Paspalum dilatatum* shoots shifted from 65° in an isolated individual to 40° for a plant in a stand of 37 plants m^-2^ (Gibson *et al*., 1992). In a mirror-image of this shift in shoot angle in response to neighbours, roots are more vertical in plant stands than for isolated individuals (Figure 1f), hence the characteristic increase in root depth with increasing plant population density (Sadras *et al*., 1989). Roots react to the presence of roots in an avoidance-type syndrome, and there is speculation about self, non-self, and kin recognition by roots (Baluška and Mancuso, 2021; Depuydt, 2014; Gruntman and Novoplansky, 2004; Hess and De Kroon, 2007). Mediated by a range of sensory traits, roots of vascular plants are central for higher-level structures that involve root–fungal networks, shared roots in clonal plants, and natural root grafts (Baluška and Mancuso, 2021).

The living parts of the environment are thus evolving phenotypes with their own genetic and environmental drivers, and their own phenotypic plasticity (Wolf *et al*., 2004). “When the environment has genes” (Wolf *et al*., 2004), the G **×** E model could be re-written as G **×** C, where C is context (Sznajder *et al*., 2010; Wolf *et al*., 2004). Context spans from cellular to ecosystem scale, *e.g.,* a gene is part of the context for another gene in intragenomic epistasis (g **×** g) at the scale of the individual or G **×** G epistasis from relationships between loci composing the genomes of different individuals in populations and communities. The mechanisms of genetic covariance are different when context is heterospecific, e.g., in plant-herbivore relations, or conspecific, e.g., plant-plant relations in crop stands, but the phenotypes are at the centre of G **×** C relations (Sznajder *et al*., 2010; Wolf *et al*., 2004).

## The causal arrow from phenotype to agronomic management: farmer’s phenotype

Phenotypic models accounting for genetic factors, environment and their interaction have been advanced for applications in human health, cognitive aptitude, ideology, and political attitudes (Ayode *et al*., 2023; Harden *et al*., 2007; McHale *et al*., 2018; Molenaar *et al*., 2013; Smith *et al*., 2011). The environment influencing the farmer’s phenotype (Box 1) includes the technological, economic and legal systems, which are unprestatable (Kauffman, 2008, 2016), and a strong biophysical component; Ballard (1962, 1965) vividly connects individual and social sense of self with the landscape transformed by climate change.

The initial spread of farming from the Levante into Central Anatolia involved the adoption of cultivars by indigenous foragers and contemporary experimentation in animal herding of local species (Baird *et al*., 2018). Communities at Boncuklu and Pinarbaşi were in broadly similar environments of the Anatolian plateau, shared technologies, and participated in the same exchange networks, but showed contrasting approaches for the exploitation of plant and animal resources in the period of approximately 8300–7800 cal BC. Both communities had almost identical foraging patterns, but the Boncuklu community adopted and sustained low-level animal husbandry and cultivation of cereals and pulses whereas the Pinarbaşi community rejected both. The reasons for these differences are unknown but correlate with contrasting social and material practices leading to two propositions: that they were distinct communities with their own distinctive identities, and that the social and symbolic significance of herding and cultivation, rather than their economic value, might have driven the earlier adoption of agronomic practices at Boncuklu (Baird *et al*., 2018).

In a context of uncertainty primarily associated with weather and market fluctuations, the causes and consequences of farmers’ risk attitudes are important. Risk attitude depends on socio-demographic characteristics, cognitive abilities, and personality attributes, and have implications for farm and crop-level decisions, technology adoption, and policy compliance (Menapace *et al*., 2013). A model of farmer’s decisions has been advanced that accounts for two traits: risk attitude and subjective belief regarding the probability of an uncertain outcome (Menapace *et al*., 2013). This model makes explicit that often individuals do not know the probability of occurrence of relevant events, and thus make decisions based upon subjective beliefs. Against this model, experiments with a relatively homogeneous sample of 313 apple farmers in northern Italy, where hail and spring frost are major risk factors, showed (i) the frequency distribution of risk attitude conformed to a power law: most farmers favour a lower payoff to avoid risk, and very few are inclined to seek a higher payoff at the expense of higher risk (Fig. 2g), and (ii) a positive association between a farmer’s level of risk aversion and their subjective belief of the probability of crop losses, which also increased with farmer’s age, previous crop losses, and exposition to outreach material (Menapace *et al*., 2013). Perceptions of risks related to climate change for growers of high-value horticultural crops were lower in the short- (*i.e.*, next season) than in the long-term (*i.e.,* 2031) and correlated with climate change beliefs after controlling for past experiences with crop losses, farming experience, numeracy, interactions with other producers, and farm characteristics (Menapace *et al*., 2015).

Differences in weed pressure between organic farms can be related to differences in farmer’s risk perception and behaviour (Riemens *et al*., 2010). A combination of surveys and measurements showed that weed pressure was higher where farmers more strongly believed that mechanical weed control compromised soil structure: weed pressure increased 10-fold from farmers who believed that mechanical weed control “never” causes soil structural damage to their counterparts who answered “often” (Fig. 2h). Differences in farmer phenotype – whatever their causes (Box 1) – influence management practices, the crop, and its environment.

Water scarcity has a two-fold effect on the crop phenotype (Fig. 2i): it compromises biological processes including plant development, nutrient uptake, growth, and resource allocation with consequences for yield, and influences farmer’s risk attitude with consequences for management decisions, which in turn affect the crop (Grassini *et al*., 2015; Pellegrini *et al*., 2022). In the US West, the rights of water users are assigned in the chronological order in which they were established (Li *et al*., 2017). Senior rights holders have priority to secure water supply, thus transferring risk to their junior counterparts. Different risk attitudes emerge from the combination of institutional and climate drivers that, in turn, influence farmer’s decisions (Li *et al*., 2017). Differences in farmer’s risk attitude associated with water availability are also apparent between irrigated and rainfed systems (Grassini *et al*., 2015) and in rainfed systems with varying frequency in the timing, intensity and duration of drought (Pellegrini *et al*., 2022). In the western US Corn Belt, the frequency of fields fertilised and protected with pesticides was lower in rainfed than in irrigated fields, and this was attributed to farmer’s reluctance to use costly inputs in inherently riskier rainfed systems (Grassini *et al*., 2015). Similarly for wheat in Argentina, the usage of fertiliser in commercial crops is lower in locations where drought is more likely, and farming is riskier (Fig. 2h).

## Conclusion

The G **×** E **×** M model of the phenotype conforms to the motif of multiple causes in cognitive maps; double-headed arrows are less prevalent. Making explicit the bidirectionality of the arrows in the G **×** E **×** M model allows to connect crop improvement and agronomy with theoretically rich fields including biological development and immunology, economics and psychology, ecology and evolution. These connections could help to narrow the gap between fast technological innovation in genotyping, phenotyping and environmental quantification, and the lagging theory of the phenotype, which is a bottleneck not only in agriculture (Sadras, 2021) but also in other technology-driven biological applications, including medicine (Nurse, 2021). Our perspective identifies, for example, two features of contemporary plant breeding that might reduce the opportunities to capture potentially valuable phenotype-to-genotype influences: nurseries managed to avoid stressful conditions and doubled haploid technologies that skip generations (Hooghvorst and Nogués, 2021).

## Acknowledgements

We thank Pedro Pellegrini for data in fig. 2i, and insights from Marta Monjardino on farmer’s risk attitude, Teresa Sadras on immunology, and Lucas Borras on plant breeding.

## Box 1.

**The farmer phenotype**

The term ‘farmer phenotype’ is rarely used in the literature on farming systems, farm management and agent modelling. A Web of Science search for the words “farmer” and “phenotype” returned 691 references mostly related to the ways farmers influence the phenotype of crops, livestock, weeds, pests and diseases, including participatory breeding where farmers select from segregating material (Annicchiarico *et al*., 2019). There is the occasional reference to anthropological studies of Neolithic farmers and hunters. Anthropologists have compared the impact of rice to wheat on cultural evolution (Talhelm and English, 2020; Talhelm *et al*., 2023). For example, the reliance on neighbours for labour and coordination of flooding and draining rice fields was used to explain why modern Chinese originating from rice growing provinces held stronger social norms than those from wheat growing regions (Talhelm and English, 2020). Tighter social norms and less-mobile relationships in rice-growing communities predicted better outcomes in the COVID-19 epidemic than in their non-rice counterparts, which were supported empirically (Talhelm *et al*., 2023).

The G **×** E **×** M framework is an example of systems thinking (Forrester, 1968) where the phenotype is treated as an emergent property of the component parts. It is an extra step to treat the manager as an emergent property of environment and genetics. The way components are treated and boundaries drawn depends on the ‘systems lens’ (Flood and Jackson, 1991; Meadows, 2008). Systems engineering with a heavy reliance on tools such as simulation modelling and operations research has been powerful to study the interactions in the context of G **×** E **×** M (Keating *et al*., 2010). Substantial thought has gone into simulating the manager’s response to the state of the system with interventions such as crop and variety choice, sowing time, and nitrogen inputs (Moore *et al*., 2014). This systems lens treats the manager as a universal rational, profit maximising, decision maker acting in isolation.

If simulation modelling represents the manager as an algorithm, precision agriculture uses algorithms and data to relieve or even replace the manager’s decision making in the same way that farm mechanisation reduced the need for manager’s physical labour (Saiz-Rubio and Rovira-Más, 2020). A reason given for the lower than hoped for adoption of variable rate fertiliser technology is the requirement for human intervention. A low-cost system that sampled the soil and then applied the fertiliser is postulated to have adoption as high as auto-steer because it eliminates the manager (Botta *et al*., 2022; Lowenberg-DeBoer, 2019).

In contrast to the lens of systems engineering, approaches embedded in the humanities, such as soft systems methodology, treat agriculture as a human activity. There is a long history of farm and farming system typology based on the structural (e.g., location, farm size) and socio-economic characteristics, and access to resources such as irrigation (Kostrowicki, 1977; Röling, 1988). More recently, differences between farmers within farming systems have led to farmer typology based on behavioural factors such as personality, worldview and interests in extrinsic financial rewards compared to intrinsic rewards of conservation (Dessart *et al*., 2019; Malek *et al*., 2019). Farmer typologies have been widely used in planning RD&E and devising policy interventions (Huber *et al*., 2024), but have not been incorporated within the G **×** E **×** M framework.

In many ways farmer typology is a synonym for farmer phenotype. Reviews of farmer typologies refer to the environment and personal styles of decision making (Bartkowski *et al*., 2022; Huber *et al*., 2024), but have not connected with the emerging research linking genetics to economic decision making or ‘genoeconomics’ (Benjamin *et al*., 2012). Behavioural genetics provides strong evidence that, while no psychological trait is 100% inheritable, all psychological traits are inheritable (Plomin *et al*., 2016). Musings on nature and nurture date back to the polymath Francis Galton in the mid-1800s, a debate that has continued for over 150 years (Pinker, 2016). This debate has been applied to risk aversion. Most of the literature on risk and decision-making refers to the environment experienced by the decision maker especially the environmental cues that frame the risky decision (Hsee and Weber, 1999; Kahneman, 2011). As discussed in the body of this paper, farmers with access to irrigation are less risk averse than farmers in rainfed systems and amongst them, risk aversion is higher in drier environments. Recently, more attention has been paid to the genetic component of risk appetite and decision making (Benjamin *et al*., 2012; De Petrillo and Rosati, 2021). Studies using standard tests for risk appetite with twins found that over half the variation has a genetic component (Nicolaou and Shane, 2019; Zyphur *et al*., 2009). A Swedish twin study showed a genetic influence on career choice with farmers ranking low on extraversion and high on risk taking (Buser *et al*., 2023). Risk appetite has been associated with hormonal responses of the neural pathways (Hogeterp *et al*., 2023; Linnér *et al*., 2019). Studies that focus on genetics do not dismiss the environment and acknowledge that phenotypes of interest to behavioural science feature complex G **×** E interactions (Driscoll, 2022; Smith *et al*., 2011).

Including management in the G **×** E framework adds complexity beyond the increase in the number of interacting components: it invites different ways of looking at the system. As pointed out by Vickers (1983) human systems are different. The agricultural economist John Dillon (1980) captured this complexity in his definition of farm management as *“the process by which resources and situations are manipulated by the farm manager in trying, with less than full information, to achieve his [or her] goals.”* The concept of farmer phenotype is relevant to Dillon’s inclusion of unique goals for each farmer and decision making under uncertainty (risk). From an anthropological perspective, Richards (1989) noted the exclusion of the farmer in G **×** E studies for small holdings. He cautioned against an over emphasis on codifying farmer knowledge and introduced the idea of farming as a performance, using the simile of a musical or theatrical performance with a script that required improvisation to perform with imperfect instruments and deal with uncertainty and surprise from nature and other performers. The metaphor of farming as a performance is also relevant to large scale, mechanised farming (Glover, 2018). The concept of the farmer phenotype influenced by the interaction of the farmer’s environment and genetic makeup contributes to an understanding of the performance.

